# The Influence of bioRχiv on *PLOS ONE*’s Peer-Review and Acceptance Time

**DOI:** 10.1101/2020.01.28.920546

**Authors:** Hiroyuki Tsunoda, Yuan Sun, Masaki Nishizawa, Xiaomin Liu, Kou Amano

## Abstract

This study examines the relation between acceptance times in preprint publishing and journal publishing. Specifically, we investigated the association between a paper’s posting time to bioRχiv, a preprints server, and journal articles’ peer-review and acceptance time for *PLOS ONE*. So far, of the total papers published in 1,626 academic journals, the average publication rate of those posted in bioRχiv is 40.67%. Meanwhile, *PLOS ONE* was the journal that published more papers. Analysis of peer-review and acceptance time of papers published in journals via preprints showed the time these papers are posted in relation to these intervals. The median of the peer-review and acceptance time of the journal submission date that was later than the date of first posting to bioRχiv was 110.00 days, and in the reverse case, it was 139.50 days. Posting to the preprint server before journal submission shows a better order than vice versa. This study provides us a good understanding of the peer-review process. It also gives us good insights into optimizing this process, which would then facilitate paper publication and knowledge dissemination.

## Introduction

Peer-review is a crucial process that ensures a submitted paper’s quality (Rowland, 2002). However, it has been widely criticized as it causes delays in the publication of new findings (Powell, 2016). Therefore, the objective of this study is to analyze the connection between the acceptance time in preprint publishing and that of journal publishing.

## Methods

First, this study investigated papers posted on bioRχiv using a web crawler that visited every page on the server and downloaded its metadata, and these papers were matched with those in the *PLOS ONE* database using the keywords of published DOIs. Second, this study investigated articles published in *PLOS ONE*. The web crawler examined *PLOS ONE* articles via bioRχiv and downloaded their received date, accepted date, and published date and then merged the metadata of bioRχiv and *PLOS ONE*. Third, peer-review and acceptance time (PT) was defined as the difference in numerical value between the accepted date and received date of the published articles. Patterns with respect to the authors’ posting process were also analyzed. The study identified two variables: preprint posted date (PP), which refers to the date when the author posted the paper to bioRχiv for the first time, and journal submission date (JS), which refers to the date when the author submitted the paper to *PLOS ONE*.

## Results and Discussion

The volume of papers posted on bioRχiv has increased rapidly every month (43,812 during November 2013–February 2019). Recently, the server has accepted more than 2,000 papers monthly, 17,818 of which were published, mostly within six months (Tsunoda et al., 2019).

**Figure 1:**
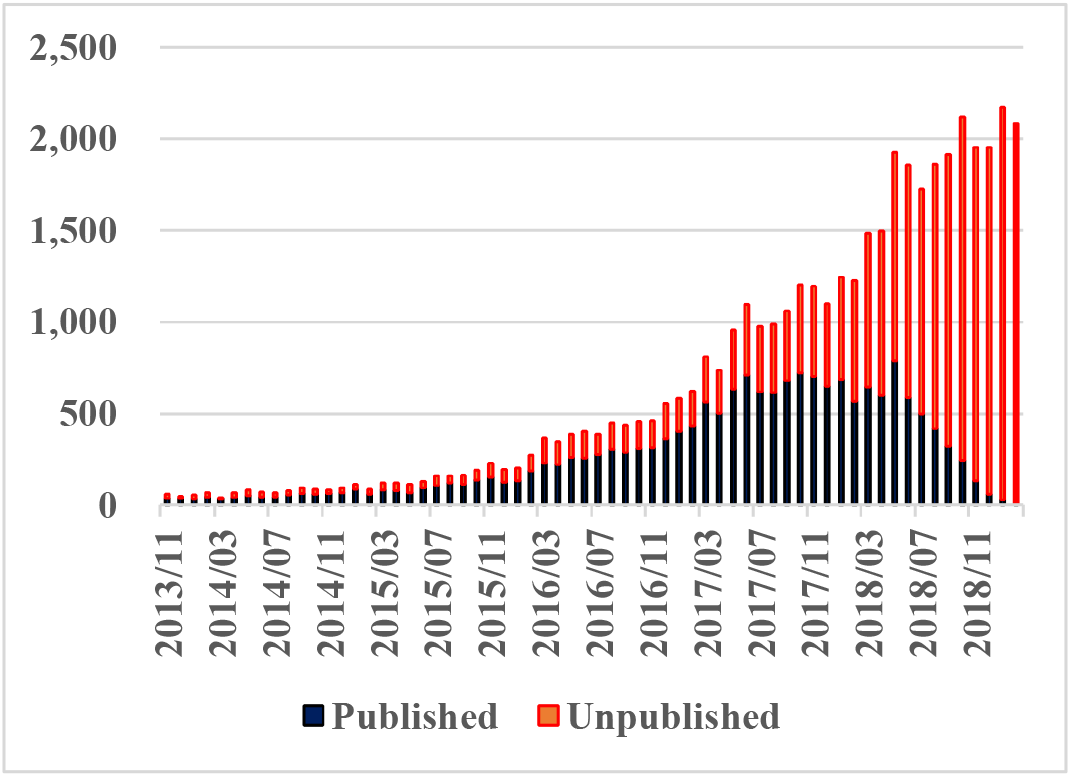
Published and Unpublished Papers Posted on bioRχiv.

These papers were published in 1,626 academic journals. Most of these papers, in terms of number and share, were in *PLOS ONE* (902 and 5.05%, respectively), followed by *Scientific Reports* (881 and 4.94%, respectively), and *eLife* (866 and 4.86%, respectively), among others.

**Figure 2:**
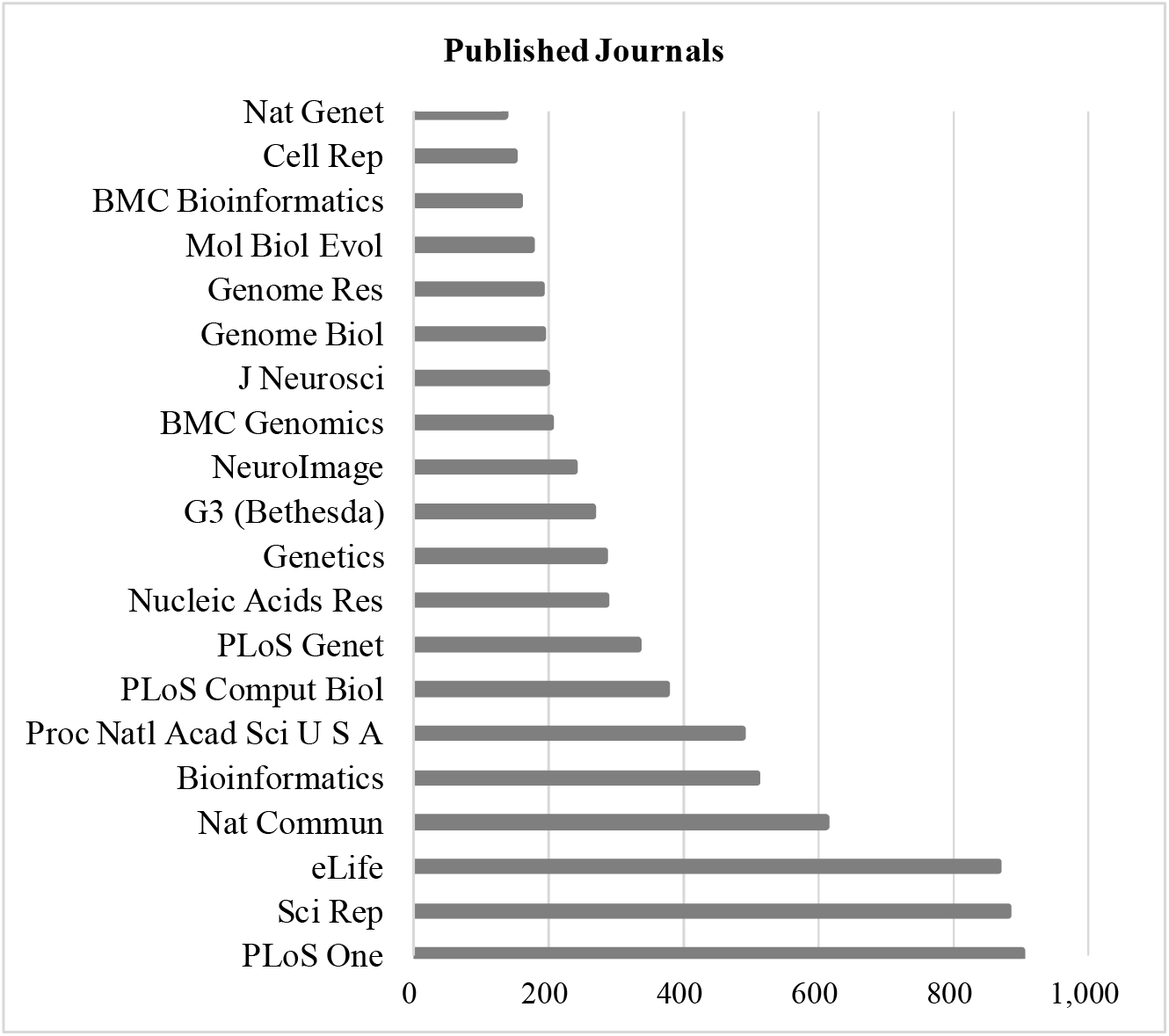
Top 20 Journals by Papers in bioRχiv.

This study focuses on *PLOS ONE* because it is a major open-access journal that has archived many biological articles, and it was bioRχiv’s top journal. PT was calculated using an equation. The shortest PT (minimum) was 7 days, the longest (maximum) was 562 days, and the middle (median) was 116 days. Quartile 1 was 82 days, and Quartile 3 was 166 days.

**Figure 3:**
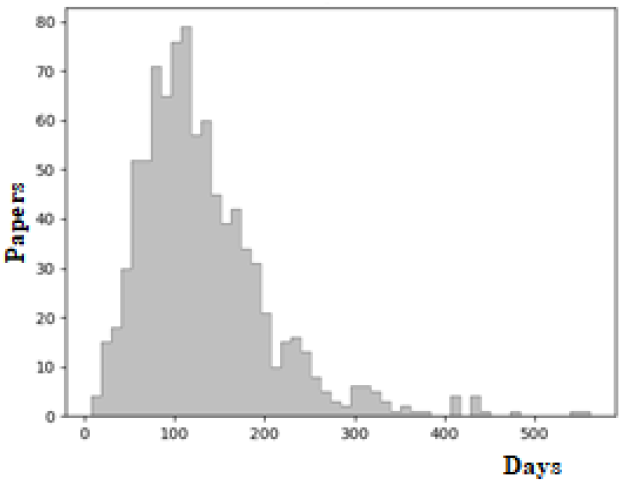
Peer Review and Acceptance Time of *PLOS ONE*.

If the difference in numerical values between preprint posted date (PP) and journal submission date (JS) was less than or equal to seven days, both dates were considered the same. This is expressed in equation (1):

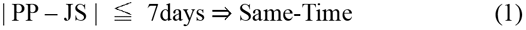

Meanwhile, if the difference in the numerical values between PP and JS is greater than seven days, PP was earlier or later than JS, as expressed in equation (2):

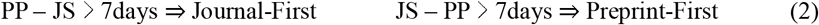

The papers’ process orders were calculated in these equations and were divided into three groups based on equation variables. Hence, group A (Same-Time) included 341 papers, group B (Preprint-First) had 333, and group C (Journal-First) consisted of 226. The PT median for Preprint-First was 110.00 days, and that for Journal-First was 139.50 days.

**Table 1:**
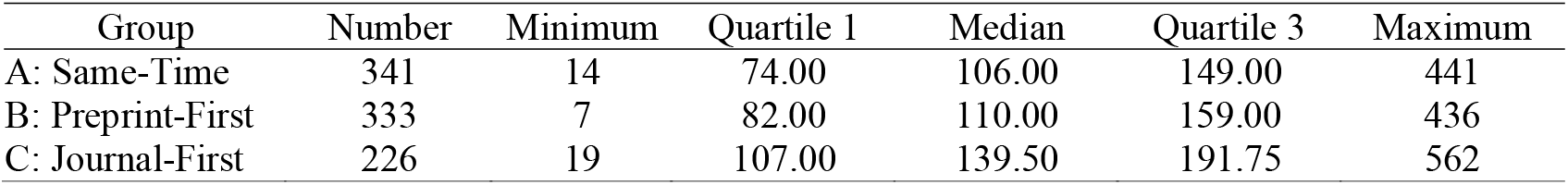
Quartiles and Median.

| Group | Number | Minimum | Quartile 1 | Median | Quartile 3 | Maximum |
| --- | --- | --- | --- | --- | --- | --- |
| A: Same-Time | 341 | 14 | 74.00 | 106.00 | 149.00 | 441 |
| B: Preprint-First | 333 | 7 | 82.00 | 110.00 | 159.00 | 436 |
| C: Journal-First | 226 | 19 | 107.00 | 139.50 | 191.75 | 562 |

**Figure 4:**
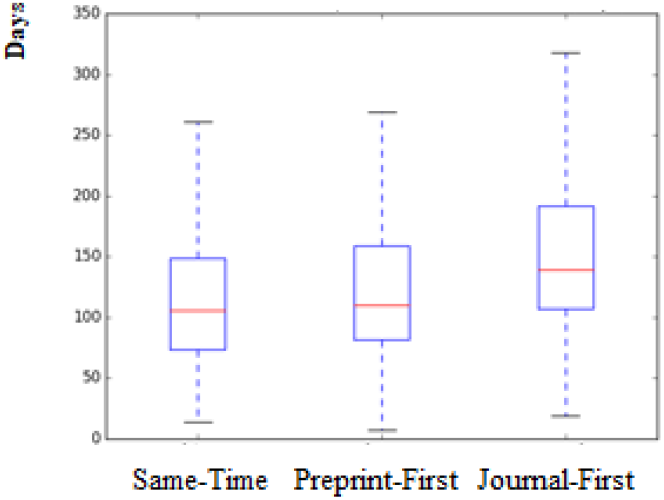
Peer Review and Acceptance Time of *PLOS ONE* (days)

**Figure 5:**
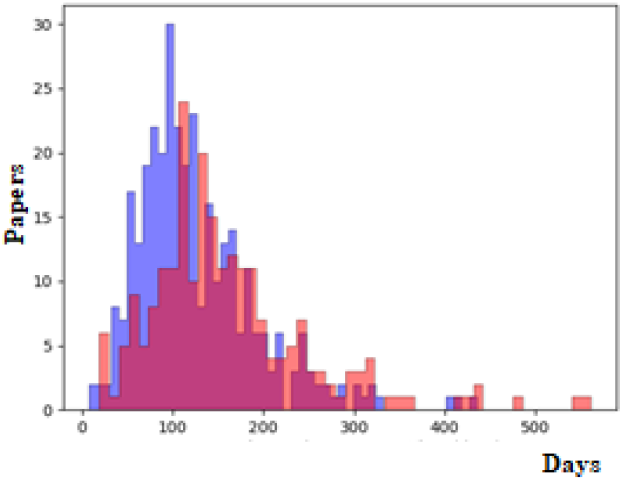
Peer Review and Acceptance Time of Group B and C. [Blue bars indicate group B, red bars indicate group C.]

The PT values for *PLOS ONE* every six months from July 2017 to June 2019 were 158, 171, 166, and 157 days (*PLOS ONE: Journal Information*, n.d.). However, the PT values in this study were shorter than normal. This could be because all articles, before being posted to bioRχiv, undergo a basic screening process for nonscientific content and checked for plagiarism. PT tends to be shorter when the preprint’s posting date is earlier than the journal’s submission date. We used the Shapiro–Wilk test to examine the normality of PT. The p-values of the Shapiro–Wilk test for Same-Time, Preprint-First, and Journal-First were about 0, which is lower than the 0.01 significance level, indicating they are not normally distributed. The median was the most suitable measure of average for PT. To compare the median of Preprint-First with that of Journal-First, the Mann–Whitney U test was used as both do not present normal distributions. The p-value here was approximately 0 and lower than the 0.01 significance level, which clearly shows that the median of Preprint-First is different from that of Journal-First. Preprint-First and Journal-First were compared because the two groups of authors displayed different tendencies. Authors who are Preprint-First could expect to receive comments and advice from scientists worldwide before submitting their papers to journals. If they were Journal-First, they cannot expect to receive such feedback.

## Conclusion

This study analyzed the PT of papers published in journals via preprints and found a relation between the time papers are posted and these intervals. If the posting date to the preprint comes before the journal submission date, PT tends to be shorter than if the order were reversed. This could be due to the basic screening process for nonscientific content and the plagiarism checks that all articles must undergo before being posted to bioRχiv. To strengthen our study, we consider to further analyze multiple different kinds of main journals, not only open access but also hybrid journals. On the other hand, in this study we only analyzed statistical data, and we need both quantitative and qualitative investigations further to make better understanding of the peer-review process, and give more insights and contribution for optimizing the peer-review process, facilitating paper publication, and disseminating knowledge.

## Acknowledgement

This work was supported by JSPS KAKENHI grant numbers JP19K12707 and JP18K11597 and ROIS NII Open Collaborative Research 2019-(19FS02).

